# Consuming Royal Jelly Causes Mosquitoes to Shift Into and Out of Their Overwintering Dormancy

**DOI:** 10.1101/2022.06.03.494749

**Authors:** Olivia E. Bianco, Aisha Abdi, Matthias S. Klein, Megan E. Meuti

## Abstract

Females of the Northern house mosquito, *Culex pipiens*, enter an overwintering dormancy, or diapause, in response to short day lengths and low environmental temperatures. Diapausing female mosquitoes feed exclusively on sugar-rich products rather than human or animal blood, thereby reducing disease transmission. During diapause, Major Royal Jelly Protein 1 (MRJP1) is upregulated in females of *Cx. pipiens*. This protein is highly abundant in royal jelly, a substance produced by honey bees (*Apis mellifera*), that is fed to future queens throughout larval development and stimulates longevity and fecundity. However, the role of MRJP1 in *Cx. pipiens* is unknown. We investigated how supplementing the diets of both diapausing and nondiapausing females of *Cx. pipiens* with royal jelly affects gene expression, egg follicle length, fat content, protein content, longevity, and metabolic profile. We found that feeding royal jelly to long day-reared females significantly reduced the egg follicle lengths of females and switched their metabolic profiles to be similar to diapausing females. In contrast, feeding royal jelly to short day-reared females significantly reduced lifespan and switched their metabolic profile to be similar nondiapausing mosquitoes. Moreover, RNAi directed against *MRJPI* significantly increased egg follicle length of short day-reared females, suggesting that these females averted diapause, although RNAi against *MRJP1* also extended the lifespan of short day-reared females. Taken together, our data show that consuming royal jelly reverses the seasonal responses of *Cx. pipiens* and that these responses are likely mediated in part by MRJP1.

**Summary Statement:** Consuming royal jelly reversed seasonal differences in physiological states, lifespan and metabolic profiles in females of the Northern house mosquito, a major vector of West Nile virus.

## 1. Introduction

The Northern house mosquito, *Culex pipiens* (L.), transmits pathogens that cause St. Louis encephalitis (Bailey et al., 1978), West Nile virus (Hamer et al., 2008), and canine heartworm (Cancrini et al., 2007) that infect millions of humans and animals each year (reviewed by Farajollahi et al., 2011). Female mosquitoes transmit these pathogens when they take a blood meal from a human or animal host (Brugman et al., 2018). However, disease transmission is not equally distributed across time (Hay et al., 2000). During the winter in temperate zones, mosquitoes enter a state of overwintering dormancy, known as diapause, where they no longer ingest blood (Bowen et al., 1988) and as a result, they no longer transmit diseases (Hay et al., 2000). Therefore, diapause in mosquitoes and other vectors has important implications for human and animal health.

Females of *Cx. pipiens* enter diapause in response to short day lengths and low environmental temperatures that act as harbingers of the approaching winter (Eldridge, 1968). Diapause allows mosquitoes to survive unfavorable winter conditions (Denlinger and Armbruster, 2014), and it involves a unique suite of behavioral, hormonal and metabolic changes (Bowen et al., 1988; Mitchell and Briegel, 1989; Robich and Denlinger, 2005). Diapause in females of *Cx. pipiens* is characterized by reproductive arrest, resulting in a decrease in the size of egg follicles as females divert energy away from reproduction (Spielman and Wong, 1973). Adult mosquitoes that enter diapause feed on sugar-rich plant nectar, causing an increase in whole body fat content (Robich & Denlinger, 2005; Sim & Denlinger, 2009; Sim et al., 2015). Several genes that regulate metabolism are differentially expressed between diapausing and nondiapausing female mosquitoes; specifically, Robich and Denlinger (2005) found that a gene associated with lipid accumulation, *fatty acid synthase*, was upregulated in diapausing females of *Cx. pipiens*, while two genes that encode enzymes related to digesting a blood meal, *trypsin* and *chymotrypsin-like* proteins, were down-regulated.

Royal jelly is produced by worker honey bees, and it is a rich source of amino acids, lipids, vitamins, and other nutrients (Colhoun and Smith, 1960). One protein in royal jelly, referred to as Major Royal Jelly Protein 1 (MRJP1), produces strong antibacterial activity and is the most abundant glycoprotein within royal jelly, constituting over 50% of total protein content and exists in a complex with the apisimin protein and 24-methylenecholesterol (Fontana et al., 2004; Ohashi et al., 1997; Tian et al., 2018). Worker bees and drones feed on royal jelly for the first 3 and 5 days of larval life respectively. Future queen bees consume royal jelly exclusively, and this induces reproductive development and extends their lifespan (Drapeau et al., 2006; Page and Peng, 2001). Surprisingly, the gene encoding MRJP1 is upregulated by the forkhead transcription factor (FOXO) and is more abundantly expressed during diapause in females of *Cx. pipiens* (Sim et al., 2015). However, the function of MRJP1 in diapausing mosquitoes is unclear, as females in diapause are not reproductively active but do live substantially longer than nondiapausing females (Spielman and Wong, 1973).

Consuming royal jelly affects the physiology of mammals, including humans, as well as fruit flies and nematodes. Integrating royal jelly into human diets can improve reproductive health, combat neurodegenerative disorders, slow aging, and promote wound-healing (reviewed by Pasupuleti et al., 2017). In rats, proteins in royal jelly can also function as antioxidants, protecting the testes of males against oxidative stress (Asadi et al., 2019). Additionally, supplementing the diets of male rams with royal jelly increases sperm motility and viability (Amini et al., 2019). Similarly, royal jelly positively impacts fertility as well as semen quality and quantity in male rabbits (El-Hanoun et al., 2014). Introducing royal jelly into the diet of *Drosophila melanogaster* (L.) extends adult lifespan in males and females, possibly by increasing antioxidant activity, and stimulates feeding behavior and fecundity in females (Xin et al., 2016). Royal jelly also extends adult lifespan in the nematode *Caenorhabditis elegans* (Maupas) (Honda et al., 2011), suggesting that royal jelly may promote longevity across a wide range of invertebrates. Moreover, Fischman et al. (2017) demonstrate that consuming royal jelly enhances the likelihood that alfalfa leaf cutting bees will enter diapause. While several studies have examined the role of royal jelly in animals, little is known about how consuming royal jelly might influence the seasonal responses and metabolic profile of mosquitoes.

Although the metabolic differences between diapausing and nondiapausing *Cx. pipiens* have not been extensively investigated, previous research has examined metabolic differences in other insects including Asian tiger mosquitoes, *Aedes albopictus* (Skuse), the flesh fly *Sarcophaga crassipalpis* (Macquart), and the parasitic wasp *Nasonia vitripennis* (Walker) (Batz and Armbruster, 2018; Michaud and Denlinger, 2007; Li et al., 2015). Not surprisingly, metabolomic studies demonstrate that diapausing mosquitoes, flesh flies, and parasitic wasps upregulate metabolites that act as cryoprotectants (Batz and Armbruster, 2018; Li et al., 2015; Michaud and Denlinger, 2007). In *N. vitripennis*, amino acids that are related to the overall metabolic pathway of glycolysis and pyruvate metabolism were differentially abundant between diapausing and nondiapausing wasps, reflecting an overall perturbation of the metabolic pathways in diapause (Li et al., 2015). Moreover, the amino acid leucine was more abundant in diapausing flesh flies while aspartate and tyrosine were more abundant in nondiapausing *S. crassipalpis* and *N. vitripennis*. However, alanine was upregulated in diapausing *S. crassipalpis* but upregulated in nondiapausing *N. vitripennis*, while glucose was upregulated in diapausing *S. crassipalpis* but upregulated in nondiapausing *N. vitripennis* (Li et al., 2015; Michaud and Denlinger, 2007). In *A. albopictus*, the monoamine neurohormones dopamine and octopamine, as well as phosphocholine and oleoyl glycine were more abundant in nondiapausing eggs compared to diapausing eggs. Preliminary data suggests that there are large-scale, global differences in the metabolic profile of diapausing and nondiapausing *Cx. pipiens* (Fig. 1), and one objective of this study is to identify specific metabolites that are differentially abundant between diapausing and nondiapausing *Cx. pipiens* and how consuming royal jelly influences the overall metabolome of short day and long day-reared mosquitoes.

**Fig. 1:**
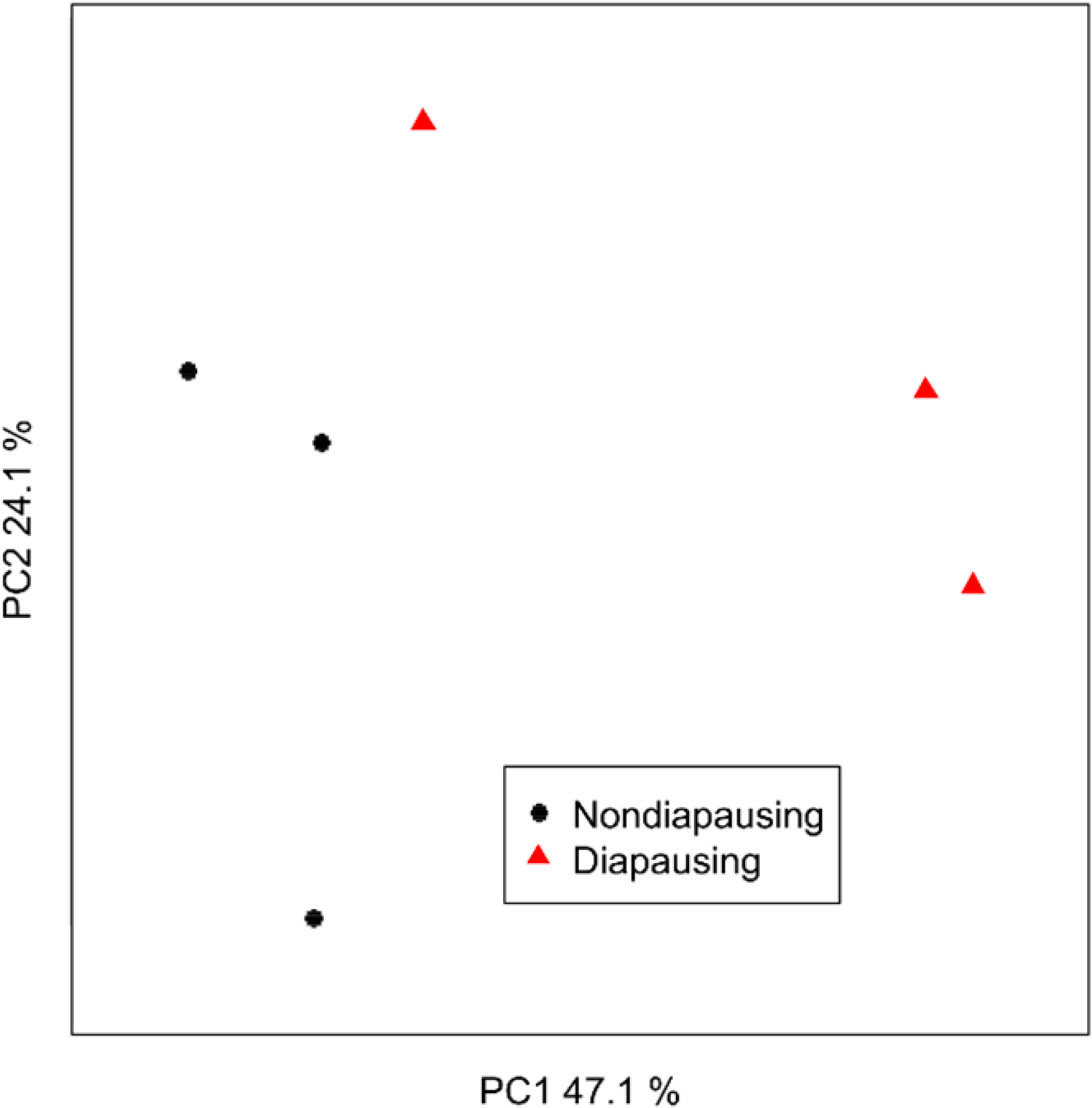
Diapausing and nondiapausing mosquitoes have different metabolic profiles. Shown is a Principal Component Analysis of whole mosquito extracts measured by 1H Nuclear Magnetic Resonance spectroscopy.

To characterize how consuming royal jelly affects seasonal responses in mosquitoes, we measured the abundance of *MRJP1* mRNA transcripts, reproductive development, lifespan, overall fat and protein content as well as the metabolic profile of long and short day-reared females of *Cx. pipiens* that had consumed royal jelly relative to sugar-water fed controls. We also assessed which, if any, of the physiological impacts of royal jelly on seasonal responses were mediated by *MRJP1* by using RNA interference (RNAi) to knock down this transcript in diapausing and nondiapausing mosquitoes. We hypothesized that mosquitoes that consumed royal jelly would enter a diapause-like state regardless of environmental conditions, characterized by small egg follicles and increased longevity, and that this would be mediated in part by an increased expression of *MRJP1*. Accordingly, we hypothesized that mosquitoes that consumed royal jelly would express a metabolic profile that was similar to that of diapausing mosquitoes that consumed sugar water. In contrast, we hypothesized that knocking down *MRJP1* with RNAi would prevent mosquitoes reared in short day, diapause-inducing conditions from entering diapause and would decrease longevity.

## 2. Materials and Methods

### 2.1. Mosquito rearing

A colony of *Culex pipiens* established in June 2013 from Columbus, Ohio (Buckeye strain) was used in this experiment. Larvae were reared at 18°C and exposed to either long day, diapause-averting conditions (photoperiod of Light:Dark 16:8 hr) or short day, diapause-inducing conditions (photoperiod of L:D 8:16 hr). Larvae were reared in plastic containers filled with reverse osmosis water, and they were fed a diet of ground fish food (Tetramin Tropical Fish Flakes) according to the procedure described by Robich and Denlinger (2005). Pupae from both the long and short-day photoperiods were divided equally into 2 cages, one of which that contained royal jelly, and the other that contained sugar water (4 treatments total; n≅150 adults/treatment). Adult mosquitoes consumed their prescribed dietary treatments *ad libitum:* sugar water (control treatment; 10% sucrose solution) or royal jelly (experimental treatment; 2 g Starkish Royal Jelly dissolved in 1.5 mL of 10% sucrose solution). One week after peak adult emergence, mosquitoes were euthanized and collected for experimental analyses.

### 2.2. Measuring the abundance of *MRJP1* mRNA

Quantitative real time PCR (qRT-PCR) was used to assess how the abundance of *MRJP1* was affected by supplementing the diet of adult mosquitoes with royal jelly. The procedure for qRT-PCR was based on Meuti et al. (2015a). We designed primers for *MRJP1* in *Cx. pipiens* (CPIJ008700-RA) using Primer3 (Forward: TGAACGATCGTCTGCTGTTC; Reverse: TCCTCCCACATGGTATCGTT; Rozen & Skaletsky, 1999). A standard curve verified that the primers met the MIQE guidelines (R^2^: 0.9885; efficiency = 108%; Bustin et al., 2009). RNA was isolated from female mosquitoes (n=5 females/biological replicate; 5 biological replicates/rearing condition and feeding treatment) one week following adult emergence using TRIzol and following the manufacturer’s instructions. Complementary DNA (cDNA) was produced using the SuperScript III kit (Invitrogen), following the manufacturer’s instructions. All qRT-PCR reactions were done in triplicate on a 96-well plate using a CFX Connect qRT-PCR machine (BioRad). Each well contained a 10 μL reaction, consisting of 5 μL of iTaq Universal SybrGreen Supermix (BioRad), 400 nM of forward and reverse primers for either *MRJP1* or our reference gene (*Ribosomal Protein 19; RpL19;* Chang and Meuti, 2020), 3.2 μL of molecular grade water, and 1 μL of cDNA. qRT-PCR reactions were completed through an initial denaturation at 94°C for 2 min, followed by 40 cycles at 94°C for 15 sec and 60°C for 1 min. The abundance of *MRJP1* transcripts was normalized to the abundance of *RpL19* using the 2^-ΔCT^ method as previously described (Chang and Meuti, 2020).

### 2.3. Assessing diapause status

To assess the diapause status of long and short day-reared females that had consumed sugar water (control) or royal jelly, we used two common markers of diapause: egg follicle length and fat content. One-week-old female mosquitoes were euthanized and dissected in a 0.9% saline solution (NaCl) using dissection needles to isolate the ovaries and egg follicles. The lengths of 10 egg follicles/female were measured and recorded at 200 times magnification using an inverted microscope (Nikon; n=20 females/treatment). The average fat content in each female mosquito was measured using a Vanillin lipid assay (Van Handel, 1985; n=8 females/treatment) that was modified to allow us to measure samples using a plate reader (Meuti et al., 2015b). The data were normalized by dividing the measured lipid content by the lean mass of the whole-body mosquito (lean mass = μg of lipid - μg wet mass).

### 2.4. Measuring protein content

As royal jelly is protein-rich, we also wanted to determine whether supplementing the diet with royal jelly affected the whole-body protein content of female mosquitoes. The protein content within individual female mosquitoes was measured using a Bradford Assay kit (BioRad) following the manufacturer’s instructions (n=8 females/treatment; Bradford, 1976). In brief, each female mosquito was weighed and then homogenized in a 2 mL microcentrifuge tube with 200 μL of a 10% ethanol solution. Samples were added in triplicate to a 96-well plate, and 250 μL of Quick Start Bradford 1X Dye Reagent (BioRad) was added to each well. The absorbance of each sample was measured using a FLUOStar Omega Microplate Reader. Measurements were normalized by dividing the protein content by the fresh mass of each female mosquito (Huck et al. 2021).

### 2.5. Evaluating longevity

To determine how dietary conditions affected the lifespan of female mosquitoes, pupae were placed in cages and were exposed to long or short-day conditions with access to sugar water or royal jelly (4 treatments total; n=100 adults/treatment). One week after adult emergence, all male mosquitoes as well as the royal jelly or sugar-water food sources were removed from each cage. The remaining females were counted and allowed constant access to water. Every week thereafter we counted and removed dead females from the bottom of the cage until no female mosquitoes remained.

### 2.6. Performing metabolomic analyses

The experimental protocol followed for tissue extraction for Nuclear Magnetic Resonance (NMR) metabolomics was based on the procedure described in Wu et al. (2008). One mosquito was weighed and placed in a 2 mL microtube with approximately 750 mg of ceramic beads (10 females per treatment group * 2 rearing conditions * 2 dietary treatments; 40 total samples). Mosquito samples were homogenized in 400 μL cold methanol and 85 μL water, and the homogenate was transferred to a separate tube without beads. Next, 400 μL chloroform and 200 μL water were added to the homogenate, which was then vortexed and centrifuged (2,000 rcf for 5 min at 4°C). The aqueous (methanol) layer was isolated and collected in a new 1.5 mL microtube before being dried in an evaporator. Deuterium oxide (heavy water), trimethylsilylpropanoic acid (TSP), and boric acid were added to the evaporated extracts and vortexed. The pH of the samples was manually adjusted to a pH of 7.4 and then transferred to 5 mm NMR tubes.

The metabolites in each mosquito sample were measured using NMR spectroscopy following the procedure detailed in Newell et al. (2018). 1D ^1^H NOESY spectra were obtained for the aqueous extracts. In addition, one ^1^H-^13^C HSQC spectrum of a pooled sample was acquired according to Gronwald et al. (2008). An Avance III HD 850 MHz spectrometer with an inverse cryoprobe and z-gradients (Bruker BioSpin, Billerica, MA) was utilized to obtain NMR measurements and resulting NMR spectra were analyzed as described in Newell et al. (2018) and Klein et al. (2011). Topspin 3.6.1 and AMIX 3.9.15 software (Bruker BioSpin, Billerica, MA) were used for preprocessing. 1D NMR spectra were binned with a bin width of 0.003 ppm using the statistical programming language R and the package mrbin (Version 1.5.0). Signals that showed large inter-spectra chemical shift differences were manually added to broader bins. Noise signals were automatically removed, and data was scaled using PQN (probabilistic quotient normalization) to correct for differences in sample mass and extraction efficacies. Each bin was then scaled to unit variance. Principal Component Analysis (PCA) models were generated to visualize metabolic data. Signals of interest were identified using public databases and identifications were validated using the acquired HSQC spectrum and measurements of pure samples.

### 2.7. Assessing the effect of knocking down *MRJP1* with RNAi

RNA interference (RNAi) was used to knock down *MRJP1* mRNA to evaluate how this protein affects the diapause status of female mosquitoes. The procedure for RNA interference was based on the protocol detailed in Meuti et al. (2015a). Double-stranded RNA (dsRNA) specific to *MRJP1* and *Beta-galactosidase (β-gal;* control; Meuti et al., 2015a) were synthesized using the Promega T7 RNAi Express Kit according to the manufacturer’s instructions. We designed primers to synthesize a 230 bp fragment of *MRJP1* (CPIJ008700-RA) in *Cx. pipiens* using Primer3 (Forward: CACCGCCAAACCGAACAAAT; Reverse: TGAGCAGCCCAAAGTACAGG; Rozen & Skaletsky, 1999), which served as the template to create dsRNA. On the day of adult emergence, 3 μg of either *β-gal* or *MRJP1* dsRNA was injected into the thorax of long and short day-reared mosquitoes. Following injection, females were placed into small plastic containers (4.62 x 6.75 x 7.19 inches) where they consumed 10% sucrose solution *ad libitium*. To confirm gene knockdown, RNA was isolated from 5 biological replicates each containing 5 whole-body, female mosquitoes two days after dsRNA injection as described above. cDNA was synthesized and qRT-PCR was conducted as described above, except that after normalizing *MRJP1* expression to the *RpL19* reference gene, *MRJP1* expression was again normalized to its expression in *β-gal*-dsRNA injected mosquitoes (Chang & Meuti, 2020). To determine how knocking down *MRJP1* affected seasonal phenotypes, the egg follicle lengths (n=20) and fat content (n=8) of females injected with dsRNA were measured ten days following injection, while the longevity of *β-gal* or *MRJP1* dsRNA-injected females (n=100) was also measured as described above.

### 2.8. Data analysis

All data analyses were conducted in R (Version 3.3.3; R Core Team, 2017). A two-way ANOVA and Tukey’s post-hoc tests were used to determine whether dietary treatment and/or photoperiod significantly affected *MRJP1* expression, egg follicle length, lipid content, and protein content in female mosquitoes. A value of alpha < 0.05 was applied to discern statistical significance. A Student’s T-test was also used to determine whether injecting *MRJP1* dsRNA effectively knocked down *MRJP1* abundance, and whether dsRNA injection significantly affected the egg follicle length or fat content of females within each rearing condition. A Kaplan-Meier survival curve and analysis were used to determine how supplementing the diet with royal jelly and knocking down *MRJP1* with RNAi affected the longevity of female mosquitoes (Therneau and Grambsch, 2000; Therneau, 2020). For analysis of NMR metabolomics data, for each spectral bin, a general linear model was created to account for the effect of diet (sugar water or royal jelly), photoperiod (long or short day-rearing conditions), and the interaction term between diet and photoperiod. Resulting p-values were corrected for multiple testing using a False Discovery Rate (FDR) of 5% (Benjamini & Hochberg, 1995).

## 3. Results

### 3.1. Measuring *MRJP1* mRNA abundance in response to rearing condition and dietary treatment

The relative abundance of *MRJP1* mRNA did not change significantly in response to dietary treatment or rearing conditions (Fig. 2A). Females reared in long day conditions that consumed royal jelly showed a slightly but not significantly higher abundance of *MRJP1* transcripts compared to those that consumed sugar water (mean relative mRNA abundance 0.0310 ± 0.00282 s.e.m.; 0.0269 ± 0.00276 s.e.m., respectively). The same trend was observed between females reared in short day conditions that consumed royal jelly (Fig. 2A; 0.0317 ± 0.00811 s.e.m.) and sugar water (0.0205 ± 0.00178 s.e.m.), but there was no significant difference. Additionally, there was no significant difference in the abundance of *MRJP1* transcripts between females reared in long day or short-day conditions.

**Fig. 2:**
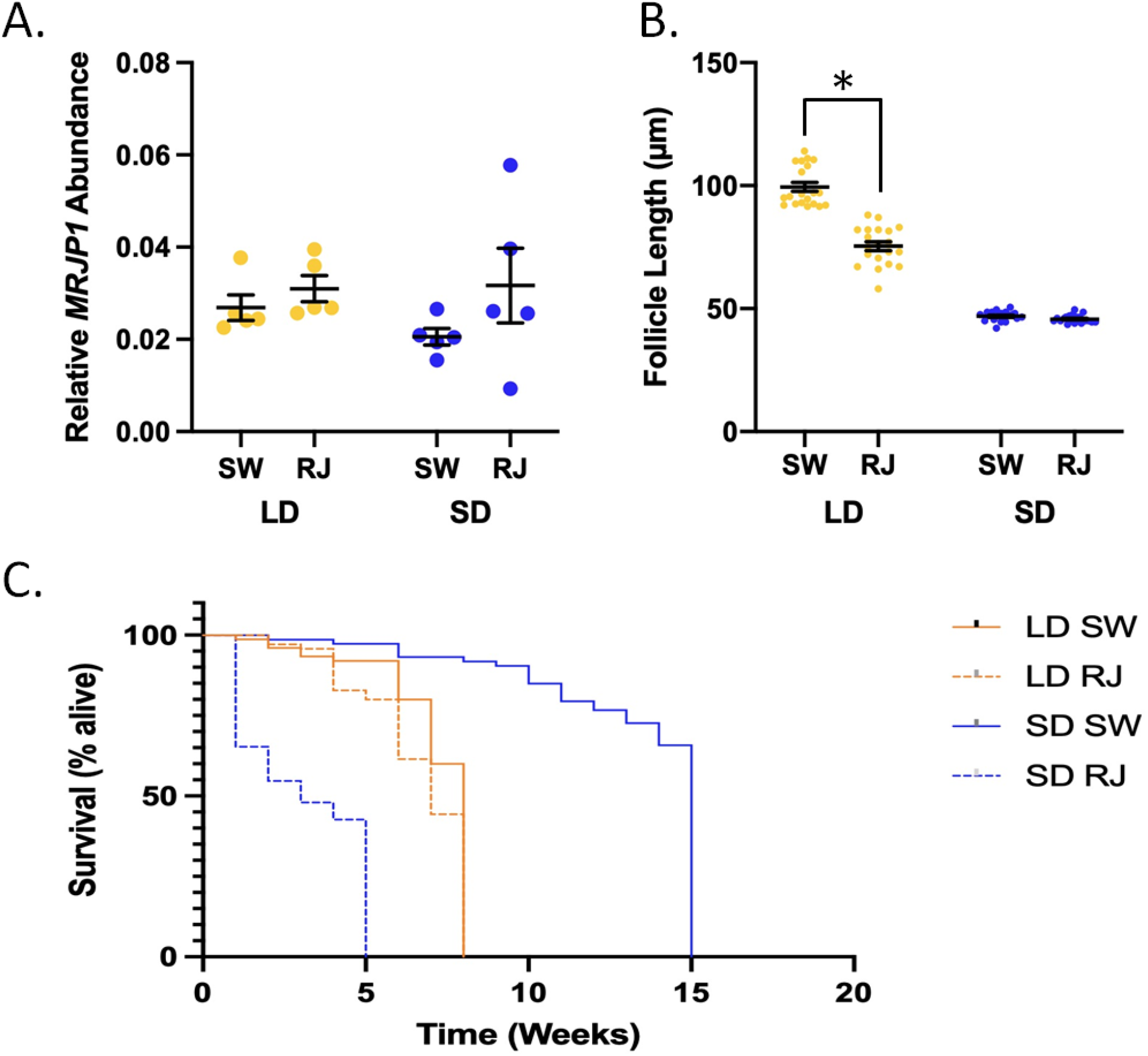
Phenotypic effects of consuming royal jelly (RJ) in mosquitoes. A.) Consuming RJ did not have any significant effect on relative *MRJP1* mRNA abundance in long day (LD) or short day (SD) reared mosquitoes. B.) Consuming RJ significantly decreases egg follicle length in long day (LD) mosquitoes. Significant difference denoted by * (p < 0.001). C.) The lifespan of female mosquitoes was significantly different between long day (LD) and short day (SD) mosquitoes. The consumption of royal jelly (RJ) by mosquitoes in both rearing conditions led to a significant decrease in lifespan. Significant differences denoted by *** (SD RJ to SD SW; p < 0.001), ** (LD SW to SD SW; p < 0.001) and * (LD RJ to LD SW; p = 0.03).

### 3.2. Assessing the effects of royal jelly on mosquito diapause status and longevity

Consuming royal jelly altered seasonal phenotypes in mosquitoes (Fig. 2B). Egg follicle length can be used to determine diapause status of female mosquitoes of *Cx. pipiens* (Robich and Denlinger, 2005; Spielman and Wong, 1973), such that an average egg follicle length of less than 75 μm indicates diapause, follicle lengths between 75 and 90 μm indicates an intermediate state, and follicles greater than 90 μm indicates non-diapause (Meuti et al., 2015b). As expected, one week after adult emergence all females reared in long-day conditions that consumed sugar water were in a clear non-diapause state (Fig. 2B; average egg follicle length of 99.5 ± 1.8 μm s.e.m.). However, 50% of long-day reared females that consumed royal jelly had egg follicles that were characteristic of being in diapause, while the remaining females were in an intermediate state, and no long day females that consumed royal jelly had egg follicle lengths that were large enough to be considered in a non-diapause state (Fig. 2B). Therefore, consuming royal jelly caused a significant decrease in the overall average egg follicle length in long day-reared females (p < 0.001), such that the average length was just above the threshold for diapause (75.4 ± 1.8 μm s.e.m.). Females reared in short-day conditions had significantly smaller egg follicles compared to those reared in long day conditions (p < 0.001). All females reared in short-day conditions were in a clear diapause state (Fig. 2B), regardless of whether they consumed sugar water (46.9 ± 0.4 μm s.e.m.) or royal jelly (45.7 ± 0.3 μm s.e.m.), and dietary treatment did not significantly impact egg follicle length of short day-reared females (p = 0.92).

We also assessed whether dietary treatment and/or photoperiod affected fat and protein content within female mosquitoes (Fig. S1), as low values for fat content indicate a non-diapause state, while higher values indicate a diapause state (Meuti et al., 2015ab). There was a non-significant increase in fat content for females reared in long day-conditions that consumed royal jelly (Fig. S1A; % lipid to lean mass 27.46 ± 4.99 % s.e.m.) compared to sugar water (16.89 ± 2.37 % s.e.m.; p = 0.21). In short day-reared females, consumption of royal jelly led to decreased levels of fat compared to sugar water, although the difference was not significant (22.19 ± 2.46 % s.e.m.; 29.48 ± 4.75 % s.e.m., respectively, p = 0.55). Females reared in long-day conditions contained roughly the same amount of protein whether they consumed royal jelly or sugar water (Fig. S1B; average protein content of 12.16 ± 0.76 μg/mg s.e.m. and 12.60 ± 1.41 μg/mg s.e.m., respectively; p = 0.99). Dietary treatment did not significantly affect the protein content of short day-reared females (average protein content in royal jelly-fed females = 10.22 ± 0.98 μg/mg s.e.m.; average protein content in sugar water-fed females 9.33 ± 1.32 μg/mg s.e.m.; p = 0.95). However, females reared in short day conditions contained significantly less protein than long day-reared females (p = 0.03).

We also investigated how the lifespan of female mosquitoes was affected by different rearing conditions and dietary treatments. There was a significant difference in lifespan between each of the four treatment groups (Fig. 2C). As expected, sugar-fed control mosquitoes reared in short-day, diapause-inducing conditions lived significantly longer than those reared in long-day, diapause-averting conditions (p < 0.001). The median survival time of long day-reared females that consumed sugar water was eight weeks of age, while 50% of sugar water-fed females survived to fifteen weeks of age in short day conditions (Fig. 2C). However, the opposite was observed with female mosquitoes that consumed royal jelly; those reared in long-day conditions lived significantly longer than those in short day-conditions (p < 0.001). The median survival time of short day-reared females that consumed royal jelly was three weeks, while 50% of long day-reared females that consumed royal jelly survived to seven weeks of age. Within long and short-day conditions, female mosquitoes that consumed sugar water lived significantly longer than those that consumed royal jelly (p = 0.03 and p < 0.001, respectively).

### 3.3. Characterizing the effects of royal jelly on the metabolomic profile of long and short day-reared mosquitoes

A Principal Component Analysis (PCA) reveals that both the photoperiodic conditions and consuming royal jelly affects the metabolism of female mosquitoes (Fig. S2). General linear models revealed that 168 spectral signals significantly changed in response to at least one of the diet, day length, and the interaction terms (Table S1). Fig. 3 shows a heatmap of all significantly changing signals. Notably, nondiapausing mosquitoes that consumed sugar water exhibit strong metabolic differences compared to diapausing mosquitoes that also consumed sugar water. However, mosquitoes reared under long-day, diapause-averting conditions switch to a “diapause” metabolic profile when consuming royal jelly. In contrast, females reared under short-day, diapause-inducing conditions switched to a “non-diapause” metabolic state when consuming royal jelly (Fig. 3). Among the significant signals, five metabolites could be unambiguously identified (Table S1). These metabolites are pimelic acid, L-alanine, asparagine, histidine, choline, and glycogen. Short day-reared, diapausing mosquitoes on that consumed sugar water and long day-reared mosquitoes that consumed royal jelly had higher levels of pimelic acid, asparagine, and choline. In contrast, short day-reared, diapausing mosquitoes that consumed sugar water and long day-reared mosquitoes that consumed royal jelly had significantly lower levels of L-alanine, histidine, and glycogen.

**Fig. 3:**
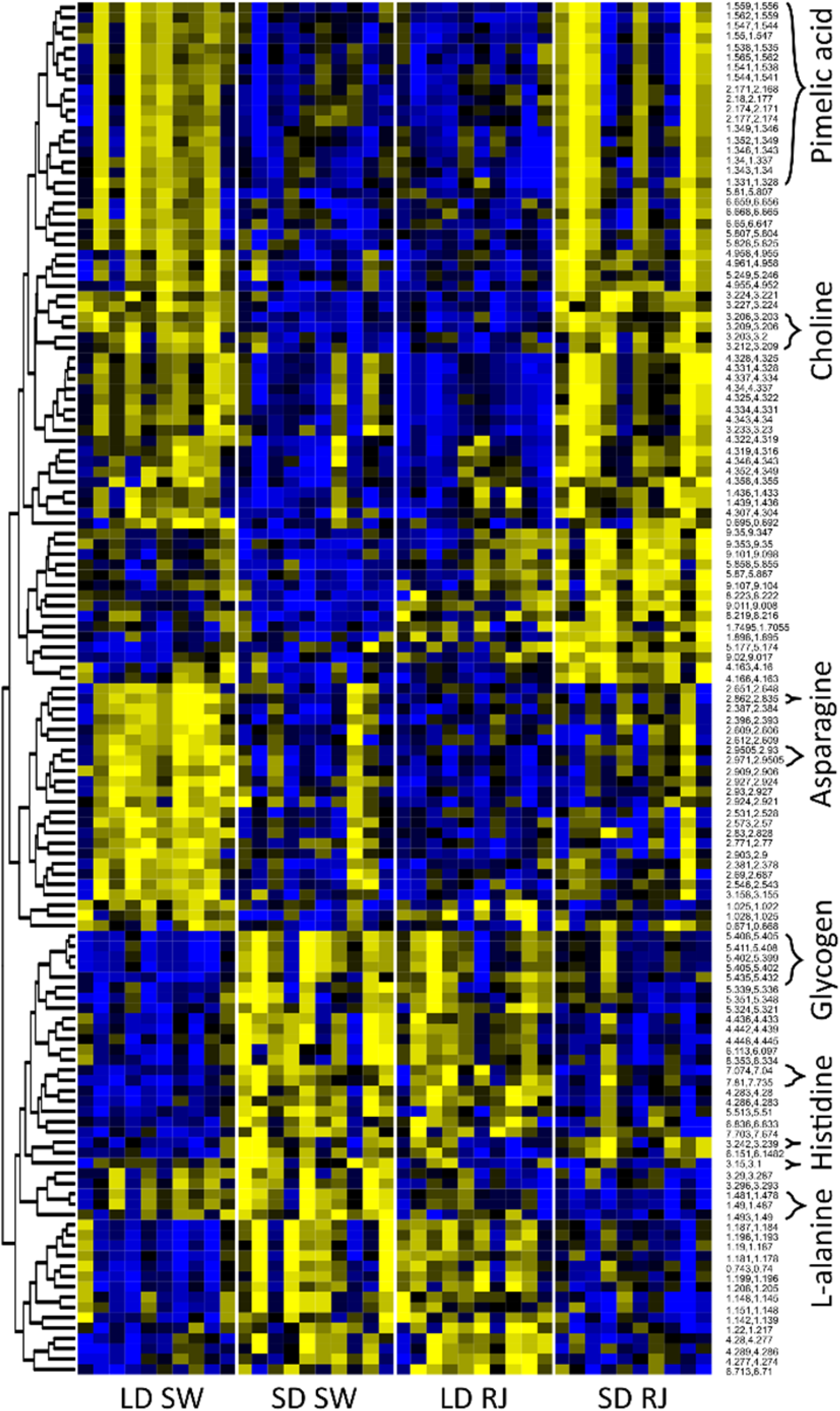
Consuming royal jelly reverses seasonal differences in the mosquito metabolome. This heat map shows regions of the NMR spectra that were significantly different between treatment groups. Signals that were unambiguously identified are labeled with the respective metabolite name. The color yellow represents metabolites that were highly abundant, while the color blue represents metabolites that were less abundant. RJ = royal jelly; SW = sugar water; SD = short-day, diapause-inducing conditions; LD = long-day, diapause-averting conditions.

### 3.4. Effects of *MRJP1* dsRNA on mosquito diapause status and longevity

A knockdown confirmation analysis was performed to determine if *MRJP1* dsRNA significantly reduced the abundance of *MRJP1* transcripts. No significant differences in mRNA abundance between *MRJP1* dsRNA and *β-gal* dsRNA-injected controls were observed in 2 independent injection trials (Fig. 4A; S3A), although there were still significant downstream effects on egg follicle length and lifespan measurements. Females reared in long day conditions treated with dsRNA for *MRJP1* had a mean relative mRNA abundance of 0.00557 ± 0.000942 s.e.m., while those treated with dsRNA for *β-gal* had 0.00532 ± 0.000947 s.e.m. (p = 0.86). For females reared in short day conditions, those treated with dsRNA for *MRJP1* had a slightly, but not significantly, lower relative mRNA abundance (Fig. 4A; 0.00470 ± 0. 0.00116 s.e.m.) compared to those treated with dsRNA for *β-gal* (0.00545 ± 0. 0.00107 s.e.m.; p = 0.25). A second round of independent injections also failed to change the levels of *MRJP1* abundance (Fig. S3A).

**Fig. 4:**
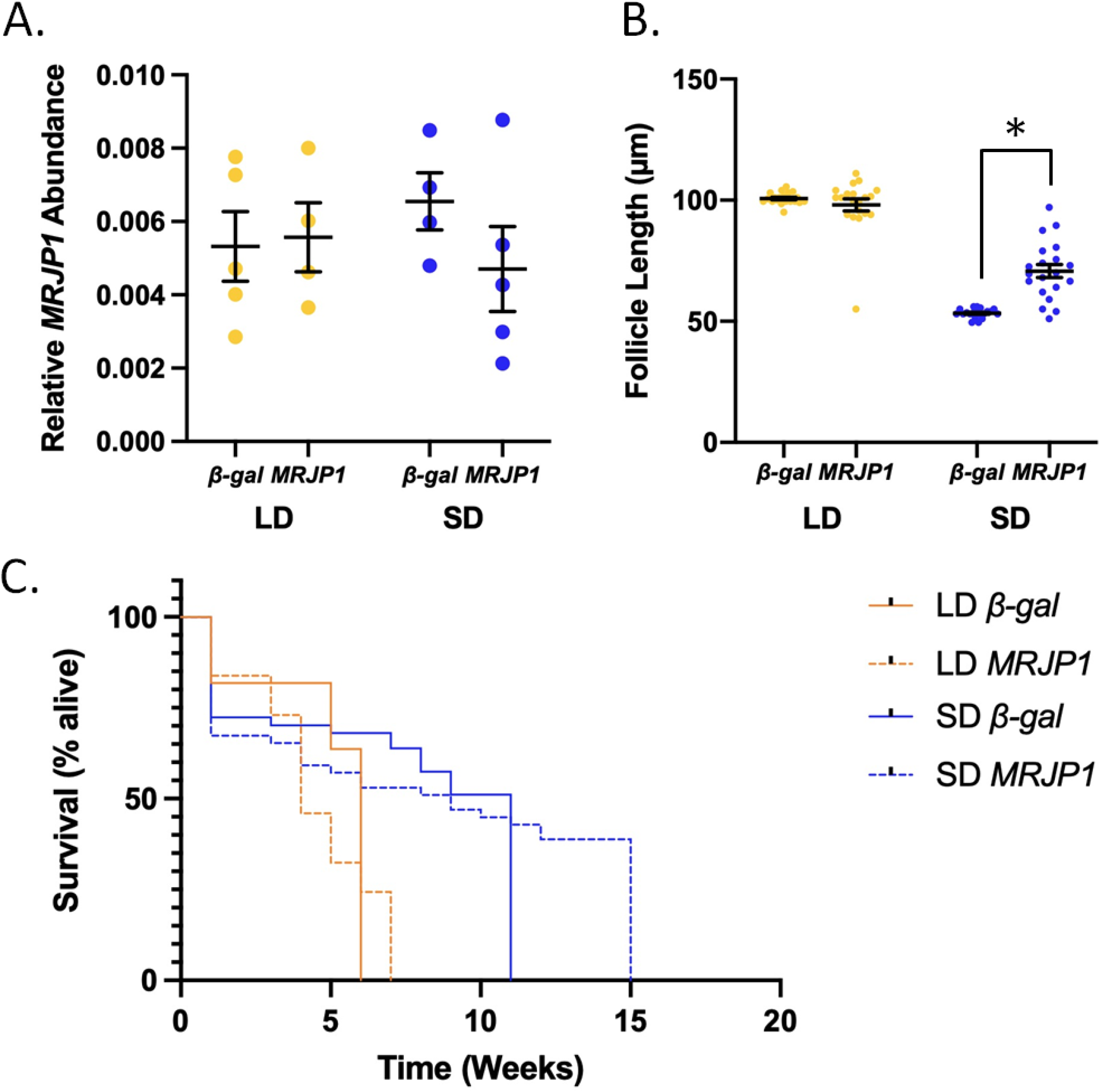
dsRNA against *MRJP1* affects seasonal phenotypes in mosquitoes. A.) Treatment with dsRNA for *β-gal* or *MRJP1* did not significantly affect relative *MRJP1* mRNA abundance in long day (LD) or short day (SD) mosquitoes. B.) dsRNA against *MRJP1* caused a significant increase in egg follicle length in short day (SD) mosquitoes. Significant difference denoted by * (p < 0.001). C.) dsRNA against *MRJP1* significantly affected the lifespan of (SD) conditions. Significant differences denoted by *** (LD *β-gal* to SD *β-gal;* p < 0.001), ** (LD *MRJP1* to SD *MRJP1;* p < 0.001) and * (SD *β-gal* to SD *MRJP1;* p = 0.0187).

All females reared in long-day conditions that were injected *β-gal* and *MRJP1* dsRNA were in a clear non-diapause state, such that the average egg follicle length of *β-gal* dsRNA-injected (100.7 ± 0.5 μm s.e.m.) and *MRJPI* dsRNA-injected mosquitoes (98.1 ± 2.5 μm s.e.m.) were not significantly different (Fig. 4B; p = 0.76). Females reared in short-day conditions that were injected with *β-gal* dsRNA were in a clear diapause state, with an average egg follicle length of 53.3 ± 0.4 μm s.e.m.. However, the average egg follicle length of females reared in short-day conditions that were injected with *MRJP1* dsRNA was 70.7 ± 2.7 μm s.e.m., and females were found to be in a mixture of diapause (70%), intermediate (25%), and non-diapause (5%) states. Overall, dsRNA directed against *MRJP1* significantly increased egg follicle length in short day-reared females (p < 0.001).

Injecting dsRNA against *MRJP1* did not significantly affect fat content (Fig. S3B). Females reared in long day conditions and treated with dsRNA for *β-gal* had an average fat content of 6.45 ± 0.78 % s.e.m., while those that were injected with *MRJP1* dsRNA had 10.97 ± 0.40 % s.e.m. (p = 0.18). Injecting *β-gal* or *MRJP1* dsRNA also had no effect on the fat content of short day-reared mosquitoes (Fig. S3B; *β-gal* dsRNA: 11.35 ± 2.46 % s.e.m.; *MRJP1* dsRNA: 11.59 ± 0.83 % s.e.m.; p = 0.99). Females reared in long day conditions and treated with dsRNA for *β-gal* had significantly less fat content compared to the other three treatment combinations (p = 0.04).

We also investigated how injection with *MRJP1* dsRNA affected the lifespan of long and short day-reared females relative to *β-gal*-injected controls (Fig. 4C). Females reared in short-day conditions that were treated with dsRNA for both *β-gal* and *MRJP1* lived significantly longer than long-day reared, dsRNA injected females (p < 0.001 and p < 0.001, respectively). For *β-gal* dsRNA-injected females, 50% of long-day reared females survived until six weeks of age while 50% of short-day reared females survived to eleven weeks of age. For *MRJP1-*injected females, 50% of long-day reared females survived until four weeks of age while 50% of short-day reared females survived until nine weeks of age. While dsRNA against *MRJP1* did not significantly affect the lifespan of females reared in long-day conditions relative to *β-gal* injected controls, females reared in short-day conditions that were treated with dsRNA for *MRJP1* had significantly longer overall lifespans than *β-gal* dsRNA-injected controls (Fig. 4C; p = 0.0187), such that the median lifespan of *MRJP1* dsRNA-injected females was 2 weeks longer than *β-gal* dsRNA-injected controls in short day conditions. (Fig. 4C).

## 4. Discussion

Our results demonstrate that consuming royal jelly reverses seasonal phenotypes in female mosquitoes. Specifically, female mosquitoes reared in long-day, diapause-averting conditions that consumed royal jelly enter a diapause-like state with small egg follicles (Fig. 2B). In contrast, consuming royal jelly significantly reduced the lifespan of females reared in short day, diapause-inducing conditions (Fig. 2C). While consuming royal jelly does not cause significant differences in fat content within long or short day-reared mosquitoes (Fig. S1A), it alters the metabolic profile of mosquitoes (Fig. 3, S2, Table S1). Specifically, long-day reared mosquitoes that consumed royal jelly were metabolically more similar to diapausing controls, while short day-reared mosquitoes that consumed royal jelly were metabolically similar to nondiapausing controls. Although it is currently not clear how royal jelly might be mediating these effects, it is likely that they are induced in large part through the action of MRJP1. This is because short day-reared female mosquitoes treated with dsRNA against *MRJP1* had significantly larger egg follicles (Fig. 4B) and lived significantly longer than *β-gal* dsRNA-injected controls (Fig. 4C).

MRJP1, a primary component of royal jelly, was upregulated in diapausing females (Sim et al., 2015). Additionally, previous studies demonstrate that consuming royal jelly increases the likelihood that alfalfa leaf cutting bees would enter diapause (Fischman et al. 2017). Therefore, we hypothesized that consuming royal jelly, and thereby artificially increasing the levels of MRJP1 within mosquitoes, would induce diapause phenotypes in long-day reared mosquitoes. Indeed, we found that consuming royal jelly significantly reduced the egg follicle lengths of long day-reared females (Fig. 2B). This finding indicates that feeding females royal jelly in conditions that typically prevent diapause causes them to arrest reproductive development. However, in contrast to our results, consuming royal jelly promotes reproductive development in honey bee queens (Page and Peng, 2001) and fruit flies (Xin et al., 2016). As other species also become more virile and fecund upon consuming royal jelly (El-Hanoun et al., 2014; Xin et al., 2016), there must be a separate, unique pathway in which MRJP1 acts in *Cx. pipiens* to confer reproductive arrest.

Reproductive arrest is not the only indicator of diapause, as diapausing females of *Cx. pipiens* also display an increase in fat content. We do observe a trend where long day-reared females that consumed royal jelly have a slightly higher fat content than females that consumed sucrose solution (Fig. S1A). However, due to high variation within our samples, the fat content is not significantly different between rearing conditions or food source in this experiment. Typically, the increase in fat content is a consequence of the feeding habits of female mosquitoes; short day-reared females gorge on nectar that is rich in sugar (Bowen et al., 1988; Robich and Denlinger, 2005; Sim and Denlinger, 2009). Although it is unclear why we did not observe significant increases in the fat content of diapausing relative to nondiapausing sugar-fed controls, we may not have observed any difference in fat content between feeding treatments because royal jelly does not necessarily have a high proportion of sugar or fat, rather it is a rich source of protein (Drapeau et al., 2006).

Considering the composition of royal jelly, we chose to examine whether protein content changed between females that consumed royal jelly relative to those who fed on sucrose (Fig. S1B). Although consuming royal jelly did not affect the protein content of long day or short day-reared mosquitoes, we do find that females reared in long-day conditions have significantly greater protein content than those reared in short-day conditions. Early in diapause, females of *Cx. pipiens* produce fewer proteins than they do upon diapause termination (Zhang et al., 2019). Furthermore, diapausing female mosquitoes are relatively inactive and take refuge in protected shelters (Eldridge, 1987), so they would not require as much protein to power their flight muscles.

In addition to determining how supplementing the diet of mosquitoes with royal jelly would affect seasonal phenotypes, we used RNAi to elucidate the functional role of MRJP1 in diapausing females of *Cx. pipiens*. Injecting *MRJP1* dsRNA in females reared in short-day, diapause-inducing conditions significant increased egg follicle length such that approximately 30% of the females sampled were considered to be in an intermediate or non-diapause state (Fig. 4B). Thus, RNAi against *MRJP1* causes short day-reared females to avert diapause, and again suggests that the gene encoding MRJP1 plays a critical role in arresting egg follicle development during diapause induction in females of *Cx. pipiens*.

RNAi against *MRJP1* does not lead to a significant change in fat content of females reared in short day conditions (Fig. S3B), although, we do observe a trend in which the long day-reared females that were injected with *MRJP1* dsRNA have a greater fat content compared to those injected with dsRNA for *β-gal*. This is a surprising trend, seeing as *MRJP1* mRNA is upregulated in diapausing mosquitoes that acquired high levels of fat (Sim et al., 2015), but we found that the fat content slightly increased when *MRJP1* was targeted with RNAi. However, this trend was not statistically significantly and likely not biologically meaningful. Overall, the female mosquitoes that were injected with *MRJP1* or *β-gal* dsRNA had lower levels of fat (Fig. S3B) than the female mosquitoes that were not injected and allowed to consume sugar water in the initial dietary experiment (Fig. S1A). We conclude that injecting the mosquitoes likely injured the mosquitoes and interfered with their ability to consume the sucrose source that was within their cage. These results, combined with the effects of consuming royal jelly, suggest that MRJP1 may not be directly involved in accumulating fat during diapause and rather that this protein regulates reproductive development and longevity.

Consuming royal jelly significantly reduced the median survival time of both long and short day-reared mosquitoes, relative to sugar-fed controls (Fig. 2C). The decrease in lifespan was most dramatic and pronounced in mosquitoes reared in short-day, diapause-inducing conditions where consuming royal jelly reduced the median lifespan by 12 weeks. This is a surprising result, seeing as royal jelly increases the lifespan of honey bees and fruit flies (Drapeau et al., 2006; Xin et al., 2016). Additionally, in short-day conditions, there was a significant difference in the longevity of female mosquitoes treated with *MRJP1* dsRNA relative to *β-gal* controls (Fig. 4C). Although *MRJP1* dsRNA-injected mosquitoes initially died sooner and had a lower median survival time than *β-gal* dsRNA injected controls, dsRNA against *MRJP1* extended the total lifespan of short day-reared mosquitoes by four weeks. Taken together, our data suggest that a factor within royal jelly, and possibly MRJP1, may reduce the lifespan of diapausing mosquitoes in a dose and time-dependent manner.

We found several metabolites that were differentially abundant between diapausing and nondiapausing mosquitoes as well as those that had consumed royal jelly (Table S1), and many of these metabolites have been associated with diapause and/or related phenotypes in other insects. Pimelic acid was upregulated during diapause and in long day-reared mosquitoes that consumed royal jelly. This compound is referred to as a survival hormone in honey bees, decreasing stress responses and suppressing lipid metabolism in honey bees that are socially isolated and that normally die quickly (Jorand et al., 1989). Although pimelic acid has not been associated with diapause in other insects, it is possible that this compound may preserve the fat content and/or promote the longevity of diapausing mosquitoes. Asparagine was also upregulated in diapausing mosquitoes, but is less abundant in diapausing larvae of the parasitoid *N. vitripennis* (Li et al., 2015). Additionally, asparagine is the most abundant free amino acid in the cotton bollworm, *Heliothis armigera* (Hübner), although its abundance does not significantly change during diapause (Boctor 1980). Concentrations of free amino acids, including asparagine, increase in the diapausing pupae of the moth, *Antheraea pernyi* (Guérin-Méneville) (Mansingh 1967). The role of asparagine in diapause is unclear, but it may confer increased cold tolerance, especially as levels of asparagine were higher in cold-acclimated granary weevils, *Sitophilus granaries* (L.), and rusty grain beetles, *Cryptolettes ferrugineus* (Stephens) (Fields et al. 1998). Additionally, we found choline is upregulated in diapausing mosquitoes. Choline is predicted to function as a cryoprotectant derived from the diet (Moos et al., 2022), which would serve a critical role in withstanding the low temperatures associated with winter.

L-alanine and histidine were downregulated in diapausing *Cx. pipiens* and long day-reared mosquitoes that consumed royal jelly. Alanine is also downregulated in diapausing *N. vitripennis* (Li et al., 2015), consistent with our findings, but is more abundant in diapausing flesh flies (Michaud and Denlinger, 2007). Alanine can serve as a cryoprotectant, its likely role in diapausing flesh flies (Michaud and Denlinger, 2007), as well as a byproduct of pyruvate metabolism. Our finding that alanine is less abundant in diapausing mosquitoes suggests that its primary role is likely in pyruvate metabolism as these mosquitoes, like diapausing wasps, are less metabolically active. Histidine is also less abundant in diapausing mosquitoes, which is consistent with findings in diapause-destined larvae of the corn earworm, *Heliocoverpa armigera* (Zhang et al., 2013). In pre-diapausing corn earworms, down-regulating histidine likely leads to lower levels of its byproduct histamine, an inhibitory neurotransmitter, that may alter the photoperiodic responses necessary for diapause induction (Zhang et al. 2013). However, histidine is upregulated in diapausing *N. vitripennis* (Li et al. 2015), and has been associated with increased cold tolerance in selected lines of *D. melanogaster* (Williams et al. 2014). As histidine combats oxidative damage (Wade and Tucker, 1998; Lemire et al. 2010), increases in histidine in *N. vitripennis* and *D. melanogaster* are likely critical for cold tolerance. It is somewhat surprising, therefore, that histidine was less abundant in diapausing *Cx. pipiens* as these females, unlike pre-diapausing *H. armigera*, have already initiated diapause and are no longer photosensitive (Sanburg and Larsen, 1973). Moreover, earlier studies have shown that diapausing *Cx. pipiens* are more cold tolerant than diapausing mosquitoes (Rinehart et al. 2006). However, previous studies have demonstrated that histidine-independent pathways that combat oxidative damage are upregulated in diapausing *Cx. pipiens* (Sim and Denlinger, 2011; King et al. 2021), which can explain why this amino acid can be downregulated in diapausing *Cx. pipiens* without compromising cold tolerance or resistance to oxidative damage.

Lastly, our findings show that glycogen was less abundant in diapausing mosquitoes and in long-day reared mosquitoes that consumed jelly. A previous study reports metabolic flux of moving dietary glucose toward glycogen in diapausing *Cx. pipiens* (King et al. 2020). However, that study did not analyze glycogen in non-diapausing animals and can thus not be directly compared to our data. Zhou and Miesfeld (2009) found that glycogen content decreases during the first weeks of diapause in *Cx. pipiens* as compared to nondiapausing animals, with a simultaneous increase in body fat. Our finding show suggest that consuming royal jelly early in diapause prevented glycogen catabolism, which likely contributed to the lower levels of body fat observed in these animals (Fig. S1A).

A quite exciting finding is that consuming royal jelly caused large scale changes in whole mosquito metabolomes that reversed the seasonal phenotypes of the mosquitoes (Figs 3 and S2). Specifically, we found that the regression coefficients of most of the significant metabolite signals is almost identical for diapausing controls and long day-reared mosquitoes that consumed royal jelly (Table S1). In contrast, the regression coefficients of the interaction between short day, diapause-inducing photoperiods and royal jelly consumption are opposite in direction and meet or exceed the sum of the diapause and royal jelly coefficients (Table S1). These results suggest that consuming royal jelly causes mosquitoes that were reared in long-day, diapause-averting conditions to display a metabolic profile that is highly similar to diapause. More impressive still, mosquitoes reared in short-day, diapause-inducing conditions that consumed royal jelly had a metabolic profile that was very similar to nondiapausing mosquitoes. These metabolic results are consistent with the phenotypes we observed, where royal jelly both induced diapause phenotypes in long day-reared mosquitoes (e.g. reduce egg follicle length) and caused short day-reared females to exhibit nondiapause phenotypes (e.g. shorten lifespan).

This the first time that a single substance has been found to reverse both long and short day seasonal phenotypes in an animal. How royal jelly switches seasonal responses is currently unclear, but we can conclude that this effect on diapause was mediated at least in part by MRJP1. This is because RNAi against *MRJP1* caused females who were reared in short-day, diapause-inducing conditions to avert diapause and develop significantly larger egg follicles. In contrast, *MRJP1* dsRNA-injected females had significantly longer lifespans. Since females in diapause do not bite humans and animals (Bowen et al. 1988), they do not transmit debilitating diseases. Future work should investigate whether it would be possible to develop control measures that use royal jelly to induce diapause in female mosquitoes during the long days of summer to reduce disease transmission (Eldridge 1987). Additionally, future work should be done to elucidate how MRJP1 and other components of royal jelly induce reproductive arrest, cause metabolic shifts and alter mosquito lifespan. Such studies will not only uncover the underpinnings of the interesting phenotypic results we observed in this study, but also lead to exciting insights on the molecular regulation of seasonal responses in other insects and mammals.

## Acknowledgements

We thank Lydia Fyie for assistance with survival and other data analyses, and Derek Huck for providing guidance and protocols on mosquito metabolomics, and additional members of the Meuti and Klein laboratories for their input and support in the design and execution of the experiments._We would also like to thank the CCIC NMR facility for their support.

## Data Availability

Upon publication, all data from the NMR metabolomics assay will be published on Dryad.

## References

Amini, S., Masoumi, R., Rostami, B., Shahir, M. H., Taghilou, P. and Arslan, H. O. (2019). Effects of supplementation of Tris-egg yolk extender with royal jelly on chilled and frozen-thawed ram semen characteristics. Cryobiology 88, 75–80.

Asadi, N., Kheradmand, A., Gholami, M., Saidi, S. H. and Mirhadi, S. A. (2019). Effect of royal jelly on testicular antioxidant enzymes activity, MDA level and spermatogenesis in rat experimental Varicocele model. Tissue Cell 57, 70–77.

Bailey, C. L., Eldridge, B. F., Hayes, D. E., Watts, D. M., Tammariello, R. F. and Dalrymple, J. M. (1978). Isolation of St. Louis encephalitis virus from overwintering *Culex pipiens* mosquitoes. Science 199, 1346–1349.

Batz, Z. A. and Armbruster, P. A. (2018). Diapause-associated changes in the lipid and metabolite profiles of the Asian tiger mosquito, Aedes albopictus. J. Exp. Biol. 221, jeb189480.

Benjamini, Y. and Hochberg, Y. (1995). Controlling the false discovery rate: a practical and powerful approach to multiple testing. J. R. Stat. Soc. B Met. 57,. 289–300.

Bowen, M. F., Davis, E. E. and Haggart, D. A. (1988). A Behavioural and Sensory Analysis of Host-Seeking Behaviour in the Diapausing Mosquito *Culex pipiens*. J. Insect Physiol. 34, 805–813.

Bradford, M. M. (1976). A rapid and sensitive method for the quantitation of microgram quantities of protein utilizing the principle of protein-dye binding. Anal. Biochem. 72, 248–254.

Brugman, V. A., Hernández-Triana, L. M., Medlock, J. M., Fooks, A. R., Carpenter, S. and Johnson, N. (2018). The role of *Culex pipiens* L. (Diptera: Culicidae) in virus transmission in Europe. Int. J. Environ. Res. Public Health 15, 389.

Bustin, S. A., Benes, V., Garson, J. A., Hellemans, J., Huggett, J., Kubista, M., Mueller, R., Nolan, T., Pfaffl, M. W., Shipley, G. L., et al. (2009). The MIQE guidelines: Minimum information for publication of quantitative real-time PCR experiments. Clin. Chem. 55, 611–622.

Cancrini, G., Scaramozzino, P., Gabrielli, S., Paolo, M. Di, Toma, L. and Romi, R. (2007). *Aedes albopictus* and *Culex pipiens* Implicated as Natural Vectors of *Dirofilaria repens* in Central Italy. J. Med. Entomol. 44, 1064–1066.

Chang, V. and Meuti, M. E. (2020). Circadian transcription factors differentially regulate features of the adult overwintering diapause in the Northern house mosquito, *Culex pipiens*. Insect Biochem. Mol. Biol. 121, 103365.

Colhoun, E. H. and Smith, M. V (1960). Neurohormonal Properties of Royal Jelly. Nature 188, 854–855.

Denlinger, D. L. and Armbruster, P. A. (2014). Mosquito Diapause. Annu. Rev. Entomol. 59, 73–93.

Drapeau, M. D., Albert, S., Kucharski, R., Prusko, C. and Maleszka, R. (2006). Evolution of the Yellow/Major Royal Jelly Protein family and the emergence of social behavior in honey bees. Genome Res. 16, 1385–1394.

El-Hanoun, A. M., Elkomy, A. E., Fares, W. A. and Shahien, E. H. (2014). Impact of royal jelly to improve reproductive performance of male rabits under hot sumer conditions. World Rabbit Sci. 22, 241–248.

Eldridge, B. F. (1968). The Effect of Temperature and Photoperiod on Blood-Feeding and Ovarian Development in Mosquitoes of the *Culex Pipiens* Complex. Am. J. Trop. Med. Hyg. 17, 133–140.

Eldridge, B. F. (1987). Diapause and Related Phenomena in *Culex* Mosquitoes: Their Relation to Arbovirus Disease Ecology. In Current Topics in Vector Research: Volume 4 (ed. Harris, K. F.), pp. 1–28. New York, NY: Springer New York.

Farajollahi, A., Fonseca, D. M., Kramer, L. D. and Marm Kilpatrick, A. (2011). “Bird biting” mosquitoes and human disease: A review of the role of *Culex pipiens* complex mosquitoes in epidemiology. Infect. Genet. Evol. 11, 1577–1585.

Fischman, B.J., Pitts-Singer, T.L. and Robinson, G.E. (2017). Nutritional regulation of phenotypic plasticity in a solitary bee (Hymenoptera: Megachilidae). Envir. Ent. 46, 1070–1079.

Fontana, R., Mendes, M. A., De Souza, B. M., Konno, K., César, L. M. M., Malaspina, O. and Palma, M. S. (2004). Jelleines: A family of antimicrobial peptides from the Royal Jelly of honeybees (*Apis mellifera*). Peptides 25, 919–928.

Hamer, G. L., Kitron, U. D., Brawn, J. D., Loss, S. R., Ruiz, M. O., Goldberg, T. L. and Walker, E. D. (2008). *Culex pipiens* (Diptera: Culicidae): A Bridge Vector of West Nile Virus to Humans. J. Med. Entomol. 45, 125–128.

Hay, S. I., Myers, M. F., Burke, D. S., Vaughn, D. W., Endy, T., Ananda, N., Shanks, G. D., Snow, R. W. and Rogers, D. J. (2000). Etiology of interepidemic periods of mosquito-borne disease. Proc. Natl. Acad. Sci. U. S. A. 97, 9335–9339.

Honda, Y., Fujita, Y., Maruyama, H., Araki, Y., Ichihara, K., Sato, A., Kojima, T., Tanaka, M., Nozawa, Y., Ito, M., et al. (2011). Lifespan-extending effects of royal jelly and its related substances on the nematode *Caenorhabditis elegans*. PLoS One 6, 1–10.

Huck, D.T., Klein, M.S. and Meuti, M.E. (2021). Determining the effects of nutrition on the reproductive physiology of male mosquitoes. J. Insect Physiol. 129, 104191.

Jorand, J. P., Bounias, M. and Chauvin, R. (1989). The “survival hormones”: Azelaic and pimelic acids, suppress the stress elicited by isolation conditions on the steroids and phospholipids of adult worker honeybees. Horm. Metab. Res. 21, 553–557.

King, B., Li, S., Liu, C., Kim, S. J. and Sim, C. (2020). Suppression of glycogen synthase expression reduces glycogen and lipid storage during mosquito overwintering diapause. J. Insect Physiol. 120, 103971.

King, B., Ikenga, A., Larsen, M. and Sim, C. (2021). Suppressed expression of oxidoreductin-like protein, Oxidor, increases follicle degeneration and decreases survival during the overwintering diapause of the mosquito *Culex pipiens*. Comp. Biochem. Phys. A. 257, 110959.

Klein, M.S., Dorn, C., Saugspier, M., Hellerbrand, C., Oefner, P.J. and Gronwald, W. (2011). Discrimination of steatosis and NASH in mice using nuclear magnetic resonance spectroscopy. Metabolomics, 7, 237–246.

Lemire, J., Milandu, Y., Auger, C., Bignucolo, A., Appanna, V.P. and Appanna, V.D. (2010). Histidine is a source of the antioxidant, α-ketoglutarate, in *Pseudomonas fluorescens* challenged by oxidative stress. FEMS Microbiol. Let. 309, 170–177.

Li, Y., Zhang, L., Chen, H., Koštál, V., Simek, P., Moos, M. and Denlinger, D. L. (2015). Shifts in metabolomic profiles of the parasitoid *Nasonia vitripennis* associated with elevated cold tolerance induced by the parasitoid’s diapause, host diapause and host diet augmented with proline. Insect Biochem. Mol. Biol. 63, 34–46.

Meuti, M. E., Stone, M., Ikeno, T. and Denlinger, D. L. (2015a). Functional circadian clock genes are essential for the overwintering diapause of the Northern house mosquito, *Culex pipiens*. J. Exp. Biol. 218, 412–422.

Meuti, M. E., Short, C. A. and Denlinger, D. L. (2015b). Mom matters: Diapause characteristics of *Culex pipiens-Culex quinquefasciatus* (Diptera: Culicidae) hybrid mosquitoes. J. Med. Entomol. 52, 131–137.

Michaud, M. R. and Denlinger, D. L. (2007). Shifts in the carbohydrate, polyol, and amino acid pools during rapid cold-hardening and diapause-associated cold-hardening in flesh flies *(Sarcophaga crassipalpis):* A metabolomic comparison. J. Comp. Physiol. B Biochem. Syst. Environ. Physiol. 177, 753–763.

Mitchell, C. J. and Briegel, H. (1989). Fate of the blood meal in force-fed, diapausing *Culex pipiens* (Diptera: Culicidae). J. Med. Entomol. 26, 332–341.

Moos, M., Korbelová, J., Štětina, T., Opekar, S., Šimek, P., Grgac, R. and Koštál, V. (2022). Cryoprotective Metabolites Are Sourced from Both External Diet and Internal Macromolecular Reserves during Metabolic Reprogramming for Freeze Tolerance in Drosophilid Fly, *Chymomyza costata*. Metabolites 12, 1–18.

Newell, C., Sabouny, R., Hittel, D., Shutt, T.E., Khan, A., Klein, M.S. and Shearer, J. (2018). Mesenchymal stem cells shift mitochondrial dynamics and enhance oxidative phosphorylation in recipient cells. Front. Physiol. 9, 1572.

Ohashi, K., Natori, S. and Kubo, T. (1997). Change in the mode of gene expression of the hypopharyngeal gland cells with an age-dependent role change of the worker honeybee *Apis mellifera* L. Eur. J. Biochem. 249, 797–802.

Page, R. E. and Peng, C. Y. S. (2001). Aging and development in social insects with emphasis on the honey bee, *Apis mellifera* L. Exp. Gerontol. 36, 695–711.

Pasupuleti, V. R., Sammugam, L., Ramesh, N. and Gan, S. H. (2017). Honey, Propolis, and Royal Jelly: A Comprehensive Review of Their Biological Actions and Health Benefits. Oxid. Med. Cell. Longev. 2017,.

R Core Team (2017). The R Project for Statistical Computing. R Found.

Rinehart, J.P., Robich, R.M. and Denlinger, D.L. (2006). Enhanced cold and desiccation tolerance in diapausing adults of Culex pipiens, and a role for Hsp70 in response to cold shock but not as a component of the diapause program. J. Med. Entomol. 43, 713–722.

Robich, R. M. and Denlinger, D. L. (2005). Diapause in the mosquito *Culex pipiens* evokes a metabolic switch from blood feeding to sugar gluttony. Proc. Natl. Acad. Sci. U. S. A. 102, 15912–15917.

Rozen, S. and Skaletsky, H. (1999). Primer3 on the WWW for General Users and for Biologist Programmers. In Bioinformatics Methods and Protocols (ed. Misener, S.) and Krawetz, S. A.), pp. 365–386. Totowa, NJ: Humana Press.

Sanburg, L.L. and Larsen, J.R. 1973. Effect of photoperiod and temperature on ovarian development in *Culex pipiens pipiens*. J. Insect Physiol. 19, 1173–1190.

Sim, C. and Denlinger, D. L. (2009). Transcription profiling and regulation of fat metabolism genes in diapausing adults of the mosquito *Culex pipiens*. Physiol. Genomics 39, 202–209.

Sim, C. and Denlinger, D.L. (2011). Catalase and superoxide dismutase-2 enhance survival and protect ovaries during overwintering diapause in the mosquito *Culex pipiens*. J. Insect Physiol. 57, 628–634.

Sim, C., Kang, D. S., Kim, S., Bai, X. and Denlinger, D. L. (2015). Identification of FOXO targets that generate diverse features of the diapause phenotype in the mosquito *Culex pipiens*. Proc. Natl. Acad. Sci. U. S. A. 112, 3811–3816.

Spielman, A. and Wong, J. (1973). Environmental control of ovarian diapause in *Culex pipiens*. Ann. Entomol. Soc. Am. 66, 905–907.

Therneau, T. M. (2020). A Package for Survival Analysis in R.

Therneau, T. M. and Grambsch, P. M. (2000). Modeling Survival Data: Extending the Cox Model. New York: Springer.

Tian, W., Li, M., Guo, H., Peng, W., Xue, X., Hu, Y., Liu, Y., Zhao, Y., Fang, X., Wang, K. and Li, X. (2018). Architecture of the native major royal jelly protein 1 oligomer. Nature Commun. 9, 1–12.

Van Handel, E. (1985). Rapid determination of total lipids in mosquitoes. J. Am. Mosq. Control Assoc. 1, 302–304.

Wade, A.M. and Tucker, H.N. (1998). Antioxidant characteristics of L-histidine. J. of Nutrit. Biochem. 9, 308–315.

Williams, C.M., Watanabe, M., Guarracino, M.R., Ferraro, M.B., Edison, A.S., Morgan, T.J., Boroujerdi, A.F. and Hahn, D.A. (2014). Cold adaptation shapes the robustness of metabolic networks in *Drosophila melanogaster*. Evolution 68, 3505–3523.

Wu, H., Southam, A.D., Hines, A. and Viant, M.R. (2008). High-throughput tissue extraction protocol for NMR-and MS-based metabolomics. Anal. Biochem. 372, 204–212.

Xin, X. X., Chen, Y., Chen, D., Xiao, F., Parnell, L. D., Zhao, J., Liu, L., Ordovas, J. M., Lai, C. Q. and Shen, L. R. (2016). Supplementation with Major Royal-Jelly Proteins Increases Lifespan, Feeding, and Fecundity in *Drosophila*. J. Agric. Food Chem. 64, 5803–5812.

Zhang, Q., Lu, Y.-X. and Xu, W.-H. (2013). Proteomic and metabolomic profiles of larval hemolymph associated with diapause in the cotton bollworm, *Helicoverpa armigera*. BMC Genomics 14, 1–13.

Zhang, C., Wei, D., Shi, G., Huang, X., Cheng, P., Liu, G., Guo, X., Liu, L., Wang, H., Miao, F., et al. (2019). Understanding the regulation of overwintering diapause molecular mechanisms in *Culex pipiens pallens* through comparative proteomics. Sci. Rep. 9, 1–12.

Zhou, G. and Miesfeld, R.L. (2009). Energy metabolism during diapause in *Culex pipiens* mosquitoes. J. Insect Physiol. 55, 40–46.

